## Supplementary information for "Rodent trapping studies as an overlooked information source for understanding endemic and novel zoonotic spillover"

### 1. Supplementary

#### 1.1. Supplementary Table 1

**Table 3**: Supplementary Table 1: Data extraction tool for studies meeting inclusion criteria

| Extraction tool | Variable | Description |
| --- | --- | --- |
| Study data |  |  |
|  | link | link to manuscript |
|  | year_publication | year of publication |
|  | title | title of manuscript |
|  | journal_name | journal |
|  | aim_1 | stated aim of study |
|  | aim_2 | stated aim of study |
|  | aim_3 | stated aim of study |
|  | first_author | first author of the study |
|  | reference_uid | DOI/ISSN/ISBN of the publication |
|  | unique_id | unique ID for the current study |
|  | metric | measurement of species presence abundance/presence |
|  | trap_types | type of rodent traps used |
|  | trapping_method | construction of the sampling grid |
|  | repeated_visit | whether there were multiple study visits to the same sites |
|  | geolocation_level | the level of geolocation reported |
|  | speciation | the level of speciation of trapped rodents |
|  | aim | aim of the study dichotomised to Ecology or Zoonotic risk |
|  | aim_detail | categorisation of study aims |
|  | species_accumulation | whether a species accumulation curve to describe trapping effort is reported |
|  | diversity_measurement | whether there is a measure of rodent species diversity reported |
|  | trapping_effort | whether pathogens are assayed |
|  | pathogen | completeness of reported trapping effort |
| Rodent data |  |  |
|  | unique_id | unique ID for the current study |
|  | year_trapping | year rodent trapping occurred (range) |
|  | month_trapping | months trapping occurred (range) |
|  | country | country trapping occurred within |
|  | region | region trapping occurred within |
|  | town_village | name of towns or villages trapping occurred within |
|  | latitude_DMS_N | latitude of trapping site in degrees minutes seconds (North) |
|  | longitude_DMS_W | longitude of trapping site in degrees minutes seconds (West) |
|  | latitude_D_N | latitude of trapping site in decimal degrees (North) |
|  | longitude_D_E | longitude of trapping site in decimal degrees (East) |
|  | UTM_coordinates | location of trapping site in UTM coordinates |
|  | habitat | habitat type of trapping site |
|  | intensity_use | the intensity of human disturbance in the trapping site |
|  | genus | reported genus of trapped rodent/small mammal species |
|  | species | reported species of trapped rodent/small mammal species |
|  | number | number of trapped individuals |
|  | trap_nights | number of trap nights reported |
|  | capture_rate | rate of capture if reported |
|  | trap_night_unit | the unit of trap night measurement |
|  | study_nights | the number of study nights completed at the trap site |
| Pathogen data |  |  |
|  | unique_id | unique ID for the current study |
|  | year_trapping | year rodent trapping occurred (range) |
|  | month | months trapping occurred (range) |
|  | country | country trapping occurred within |
|  | region | region trapping occurred within |
|  | town_village | name of towns or villages trapping occurred within |
|  | habitat | habitat type of trapping site |
|  | genus | reported genus of trapped rodent/small mammal species |
|  | species | reported species of trapped rodent/small mammal species |
|  | pathogen_x | pathogens tested for, 1-7 possible columns |
|  | latitude_DMS_N | latitude of trapping site in degrees minutes seconds (North) |
|  | longitude_DMS_W | longitude of trapping site in degrees minutes seconds (West) |
|  | latitude_D_N | latitude of trapping site in decimal degrees (North) |
|  | longitude_D_E | longitude of trapping site in decimal degrees (East) |
|  | UTM_coordinates | location of trapping site in UTM coordinates |
|  | path_x_tested | number of individuals assayed for the corresponding pathogen, 1-7 possible columns |
|  | pcr_x_positive | number of individuals PCR positive for the corresponding pathogen, 1-7 possible columns |
|  | ab_ag_x_positive | number of individuals with positive serological assays for the corresponding pathogen, 1-7 possible columns |
|  | culture_x_positive | number of individuals culture positive for the corresponding pathogen, 1-7 possible columns |
|  | histo_x_positive | number of individuals histologically/histopathologically positive for the corresponding pathogen, 1-7 possible columns |

#### 1.2. Supplementary Table 2

Supplementary Table 2: Included studies

| Year publication | Author | Title | Journal/Publication |
| --- | --- | --- | --- |
| 1974 | D. C. D. Happold | The small rodents of the forest-savanna-farmland association near Ibadan, Nigeria, with observations on reproduction biology | Revue de Zoologie et de Botanique Africaines |
| 1974 | Thomas Monath | Lassa virus isolation from Mastomys natalensis rodents during an epidemic in Sierra Leone | Science |
| 1975 | Herta Wulff | Recent isolations of Lassa virus from Nigerian rodents | Bulletin of the World Health Organisation |
| 1975 | Malcolm Coe | Mammalian ecological studies on Mount Nimba, Liberia. | Mammalia |
| 1977 | D. C. D. Happold | A population study on small rodents in the tropical rain forest of Nigeria | La Terre et la Vie |
| 1977 | Sonia Jeffrey | Rodent Ecology and Land Use in Western Ghana | Journal of Applied Ecology |
| 1979 | J P Dedet | Isolation of Leishmania major from Mastomys erythroleucus and Tatera gambiana in Senegal (West Africa). | Annals of Tropical Medicine and Parasitology |
| 1982 | A Diallo | Bacteriological survey of leptospirosis in Zaria, Nigeria. | Tropical and geographical medicine |
| 1983 | C Robbins | Mastomys (rodentia: muridae) species distinguished by hemoglobin pattern differences. | The American journal of tropical medicine and hygiene |
| 1987 | Joseph McCormick | A prospective study of the epidemiology and ecology of Lassa fever. | Journal of Infectious Diseases |
| 1988 | J. Iyawe | Distribution of small rodents and shrews in a lowland rain forest zone of Nigeria, with observations on their reproductive biology | African Journal of Ecology |
| 1991 | J Trape | Tick-borne Borreliosis in West Africa | The Lancet |
| 1992 | Brian Mahy | Maintenance support of a field station in Sierra Leone, West Africa | US Army Medical Research and Development Command |
| 1993 | Laurent Granjon | Social structure in synanthropic populations of a murid rodent Mastomys natalensis in Senegal | Acta Theriologica |
| 1994 | Bruno Godeluck | A longitudinal survey of Borrelia crocidurae prevalence in rodents and insectivores in Senegal. | American Journal of Tropical Medicine and Hygiene |
| 1994 | G Diatta | A comparative study of three methods of detection of Borrelia crocidurae in wild rodents in Senegal | Transactions of the Royal Society of Tropical Medicine and Hygiene |
| 1995 | E Ikeh | Mastomys natalensis and Tatera gambiana as probable reservoirs of human cutaneous leishmaniasis in Nigeria. | Transactions of the Royal Society of Tropical Medicine and Hygiene |
| 1997 | C Mafiana | Gastrointestinal helminth parasites of the black rat (Rattus rattus) in Abeokuta, southwest Nigeria. | Journal of Helminthology |
| 1999 | Holger Meinig | Notes on the mammal fauna of the southern part of the Republic of Mali, West Africa. | Bonn Zoological Bulletin |
| 1999 | Jan Decher | Diversity and structure of terrestrial small mammal communities in different vegetation types on the Accra Plains of Ghana | Journal of Zoology |
| 2000 | Adrian Barnett | Ecology of rodent communities in agricultural habitats in eastern Sierra Leone: Cocoa groves as forest refugia | Tropical Ecology |
| 2000 | J Duplantier | Rodents as reservoir hosts in the transmission of Schistosoma mansoni in Richard-Toll, Senegal, West Africa | Journal of Helminthology |
| 2000 | James Ryan | Mammal fauna of the Muni-Pomadze Ramsar site, Ghana. | Biodiversity and Conservation |
| 2001 | Austin Demby | Lassa Fever in Guinea: {II}. Distribution and Prevalence of Lassa Virus Infection in Small Mammals | Vector-Borne and Zoonotic Diseases |
| 2001 | Khalilou Ba | Preliminary study on some rodents of southern Mauritania as rÃ©servoir of human pathogenic viruses | African Small Mammals |
| 2002 | Gauthier Dobigny | A cytotaxonomic survey of Rodents from Niger: implications for systematics, biodiversity and biogeography | Mammalia |
| 2002 | Laurent Granjon | The small mammal community of a coastal site of south-west Mauritania | African Journal of Ecology |
| 2003 | D Attuquayefio | A study of bushfires in a Ghanaian coastal wetland. Impact on small mammals | African Journal of Applied Ecology |
| 2003 | Laurent Granjon | The importance of cytotaxonomy in understanding the biogeography of African rodents: Lake Chad murids as an example | Mammal Review |
| 2004 | B. Sicard | Effects of climate and local aridity on the latitudinal and habitat distribution of Arvicanthis niloticus and Arvicanthis ansorgei (Rodentia, Murinae) in Mali | Journal of Biogeography |
| 2004 | Jan Decher | A rapid survey of terrestrial small mammals (shrews and rodents) of the Foret Classee du Pic de Fon, Guinea. | Rapid Assessment Program |
| 2004 | Sara Churchfield | First results on the feeding ecology of sympatric shrews (Insectivora: Soricidae) in the Tai National Park, Ivory Coast | Acta Theriologica |
| 2005 | Ara Monadjem |  | Conservation International |
| 2005 | D Attuquayefio | Preliminary biodiversity assessment (herpetofauna and mammals) of a coastal wetland in the Volta region, Ghana | Ghana Journal of Science |
| 2005 | Francesco Angelici | Patterns of specific diversity and population size in small mammals from arboreal and ground-dwelling guilds of a forest area in southern Nigeria | Journal of Zoology |
| 2005 | Jan Decher | Rapid assessment of Small Mammals at Draw River, {BoiTano}, and Krokosua Hills | Conservation International |
| 2005 | Laurent Granjon | Population dynamics of the multimammate rat Mastomys huberti in an annually flooded agricultural region of central Mali | Journal of Mammology |
| 2006 | Natalie Weber | A Rapid Survey of Small Mammals from the Atewa Range Forest Reserve, Eastern Region, Ghana | Conservation International |
| 2006 | Patrick Barriere | Rapid Survey of the Small Mammals of Ajenjua Bepo and Mamang River Forest Reserves, Ghana | Conservation International |
| 2006 | Ryan Norris | A rapid biological assessment of three classified forests in southeastern Guinea | RAP Bulletin of Biological Assessment 40 |
| 2007 | Joseph Fair | Lassa Virus-Infected Rodents in Refugee Camps in Guinea: A Looming Threat to Public Health in a Politically Unstable Region | Vector-Borne and Zoonotic Diseases |
| 2008 | Ayodeji Olayemi | Diversity and distribution of murid rodent populations between forest and derived savanna sites within south western Nigeria. | Biodiversity and Conservation |
| 2008 | D Attuquayefio | Biodiversity assessment (rodents and avifauna) of five forest reserves in the Brong-Ahafo Region, Ghana | Ghana Journal of Science |
| 2008 | G Raczniak | Cutaneous leishmaniasis in the Volta district of Ghana: An uncertain reservoir for focal disease outbreak | The Libyan Journal of Infectious Diseases |
| 2008 | Laurent Crespin | Annual flooding, survival and recruitment in a rodent population from the Niger River plain in Mali | Journal of Tropical Ecology |
| 2009 | Christiane Denys | New data on the taxonomy and distribution of Rodentia (Mammalia) from the western and coastal regions of Guinea West Africa | Italian Journal of Zoology |
| 2009 | Elisabeth Fichet-Calvet | Diversity and dynamics in a community of small mammals in coastal Guinea, West Africa | Belgian Journal of Zoology |
| 2009 | Ivoke Njoku | Studies on the seasonal variations and prevalence of helminth fauna of the black rat, Rattus rattus (L) (Rodentia: Muridae) from different microhabitats in Nsukka, Nigeria. | Animal Research International |
| 2010 | Adam Konecny | Indications of higher diversity and abundance of small rodents in human-influenced Sudanian savannah than in the Niokolo Koba National Park (Senegal). | African Journal of Ecology |
| 2010 | David Safronetz | Detection of Lassa Virus, Mali | Emerging Infectious Diseases |
| 2010 | Edward Omudu | A survey of rats trapped in residential apartments and their ectoparasites in Makurdi, Nigeria. | Research Journal of Agriculture and Biological Sciences |
| 2010 | Elisabeth Fichet-Calvet | Diversity, dynamics and reproduction in a community of small mammals in Upper Guinea, with emphasis on pygmy mice ecology | African Journal of Ecology |
| 2010 | Gauthier Dobigny | Molecular survey of rodent-borne Trypanosoma in Niger with special emphasis on T. lewisi imported by invasive black rats | Acta Tropica |
| 2010 | Jan Decher | Small mammal survey in the upper Seli River valley, Sierra Leone | Mammalia |
| 2010 | Lies Durnez | Terrestrial Small Mammals as Reservoirs of Mycobacterium ulcerans in Benin | Applied and Environmental Microbiology |
| 2010 | Mary Reynolds | A Silent Enzootic of an Orthopoxvirus in Ghana, West Africa: Evidence for Multi-Species Involvement in the Absence of Widespread Human Disease | American Journal of Tropical Medicine and Hygiene |
| 2010 | R Sall-Drame | Variation in cestode assemblages of Mastomys and Arvicanthis species (Rodents: Muridae) from Lake Retba in Western Senegal. | Journal of Parasitology |
| 2010 | Violaine Nicolas | Terrestrial small mammal diversity and abundance in central Benin: comparison between habitats, with conservation implications | African Journal of Ecology |
| 2011 | David Coulibaly-N'golo | Novel Arenavirus Sequences in Hylomyscus sp. and Mus (Nannomys) setulosus from Cote d'Ivoire: Implications for Evolution of Arenaviruses in Africa | PLOS One |
| 2011 | Karmidine Hima | Extensive Robertsonian polymorphism in the African rodent Gerbillus nigeriae: geographic aspects and meiotic data | Journal of Zoology |
| 2011 | Laurent Granjon | Guinean biodiversity at the edge: Rodents in forest patches of southern Mali | Mammalian Biology |
| 2011 | M. Thiam | Capacity for water conservation in invasive (Gerbillus nigeriae) and declining rodents (Taterillus pygargus and Taterillus gracilis) that exhibit climate-induced distribution changes in Senegal | Journal of Arid Environments |
| 2012 | Amawulu Ebenezer | Effects of Urbanization and Agricultural Expansion on the Upsurge of Wild Rats (Rattus rattus) in Yenagoa Metropolis of Bayelsa State, Nigeria | Research Journal of Applied Sciences, Engineering and Technology |
| 2012 | Christiane Denys | On a new species of Dendromus (Rodentia, Nesomyidae) from Mount Nimba, Guinea. | Mammalia |
| 2012 | Khalilou Ba | Ecology of a typical West African Sudanian savannah rodent community. | African Journal of Ecology |
| 2012 | Laurent Crespin | Demographic aspects of the island syndrome in two Afrotropical Mastomys rodent species | Acta Oecologica |
| 2012 | Tom Schwan | Endemic Foci of the Tick-Borne Relapsing Fever Spirochete Borrelia crocidurae in Mali, West Africa, and the Potential for Human Infection | PLOS NTD |
| 2013 | Adam Konecny | Invasion genetics of the introduced black rat (Rattus rattus) in Senegal, West Africa. | Molecular Ecology |
| 2013 | Blaise Kadjo | Assessment of terrestrial small mammals and a record of the critically endangered shrew Crocidura wimmeri in Banco National Park (Cote d'Ivoire) | Mammalia |
| 2013 | Gualbert Houemenou | Leptospira spp. Prevalence in Small Mammal Populations in Cotonou, Benin | ISRN Epidemiology |
| 2013 | Jean-Francois Trape | The epidemiology and geographic distribution of relapsing fever borreliosis in West and North Africa, with a review of the Ornithodoros erraticus complex (Acari: Ixodida). | PLOS One |
| 2013 | Joshua Kamani | Prevalence and diversity of Bartonella species in commensal rodents and ectoparasites from Nigeria, West Africa. | PLOS NTD |
| 2013 | Karl Kronmann | Two Novel Arenaviruses Detected in Pygmy Mice, Ghana | Emerging Infectious Diseases |
| 2013 | Reuben Garshong | Effect of Habitat Change through Infrastructural Development on Small Mammal Diversity and Abundance on the Legon Campus of the University of Ghana. | West African Journal of Applied Ecology |
| 2014 | Benjamin Ofori | Preliminary checklist and aspects of the ecology of small mammals at the University of Ghana Botanical Garden, Accra Plains, Ghana | Journal of Biodiversity and Environmental Sciences |
| 2014 | Elisabeth Fichet-Calvet | Lassa Serology in Natural Populations of Rodents and Horizontal Transmission | Vector-Borne and Zoonotic Diseases |
| 2014 | Madougou Garba | Spatial Segregation between Invasive and Native Commensal Rodents in an Urban Environment: A Case Study in Niamey, Niger | PLOS One |
| 2015 | Charles Narh | Source Tracking Mycobacterium ulcerans Infections in the Ashanti Region, Ghana | PLOS NTD |
| 2015 | Christelle Dassi | Detection of Mycobacterium ulcerans in Mastomys natalensis and Potential Transmission in Buruli ulcer Endemic Areas in CÃ´te d'Ivoire | Mycobacterial Diseases |
| 2015 | Georges Diatta | Borrelia infection in small mammals in West Africa and its relationship with tick occurrence inside burrows | Acta Tropica |
| 2015 | Pilar Foronda | Serological survey of antibodies to Toxoplasma gondii and Coxiella burnetii in rodents in north-western African islands (Canary Islands and Cape Verde). | Onderstepoort Journal of Veterinary Research |
| 2015 | R Mol | Small terrestrial mammal and amphibian survey BoÃ© region, Guinea-Bissau. | Silvavir Forest Consultants |
| 2015 | Thomasz Leski | Sequence variability and geographic distribution of Lassa virus, Sierra Leone. | Emerging Infectious Diseases |
| 2016 | Ayodeji Olayemi | Arenavirus Diversity and Phylogeography of Mastomys natalensis Rodents, Nigeria | Emerging Infectious Diseases |
| 2016 | Ayodeji Olayemi | New Hosts of The Lassa Virus | Scientific Reports |
| 2016 | Benjamin Ofori | Spatio-temporal variation in small mammal species richness, relative abundance and body mass reveal changes in a coastal wetland ecosystem in Ghana | Environmental Monitoring and Assessment |
| 2017 | Alexis Ribas | Whipworm diversity in West African rodents: a molecular approach and the description of Trichuris duplantieri n. sp (Nematoda: Trichuridae) | Parasitological Research |
| 2017 | C Lippens | Genetic structure and invasion history of the house mouse (Mus musculus domesticus) in Senegal, West Africa: a legacy of colonial and contemporary times | Heredity |
| 2017 | Christophe Diagne | Ecological and sanitary impacts of bacterial communities associated to biological invasions in African commensal rodent communities | Scientific Reports |
| 2017 | Christophe Diagne | Serological Survey of Zoonotic Viruses in Invasive and Native Commensal Rodents in Senegal, West Africa | Vector-Borne and Zoonotic Diseases |
| 2017 | Daniel Attuquayefio | Impact of mining and forest regeneration on small mammal biodiversity in the Western Region of Ghana | Environmental Monitoring and Assessment |
| 2018 | Ayodeji Olayemi | Widespread arenavirus occurrence and seroprevalence in small mammals, Nigeria | Parasites and Vectors |
| 2018 | Benjamin Ofori | Urban green area provides refuge for native small mammal biodiversity in a rapidly expanding city in Ghana | Environmental Monitoring and Assessment |
| 2018 | Carine Brouat | Seroprevalence of Toxoplasma gondii in commensal rodents sampled across Senegal, West Africa. | Parasite |
| 2018 | Isaac Clement | Endoparasites of Small Mammals in Edo State, Nigeria: Public Health Implications | The Korean Journal of Parasitology |
| 2018 | Joshua Kamani | Prevalence of Hepatozoon and Sarcocystis spp. in rodents and their ectoparasites in Nigeria | Acta Tropica |
| 2018 | Katharina Schaufler | Clinically Relevant {ESBL}-Producing K. pneumoniae {ST}307 and E. coli {ST}38 in an Urban West African Rat Population | Frontiers in Microbiology |
| 2018 | Kouame Akpatou | Terrestrial small mammal diversity and abundance in TaÃ¯ National Park, CÃ´te d'Ivoire | Nature Conservation Research |
| 2018 | Lokman Galal | Diversity of Toxoplasma gondii strains shaped by commensal communities of small mammals. | International Journal for Parasitology |
| 2018 | Marien Joachim | Movement Patterns of Small Rodents in Lassa Fever-Endemic Villages in Guinea | EcoHealth |
| 2018 | Stefano Catalano | Rodents of Senegal and their role as intermediate hosts of Hydatigera spp. (Cestoda: Taeniidae). | Parasitology |
| 2019 | Agnes Yadouleton | Lassa Virus in Pygmy Mice, Benin, 2016â€“2017 | Emerging Infectious Diseases |
| 2019 | Gualbert Houemenou | Pathogenic Leptospira in Commensal Small Mammals from the Extensively Urbanized Coastal Benin | Urban Science |
| 2019 | Joachim Marien | Evaluation of rodent control to fight Lassa fever based on field data and mathematical modelling. | Emerging Microbes and Infections |
| 2019 | Karmidine Hima | Native and Invasive Small Mammals in Urban Habitats along the Commercial Axis Connecting Benin and Niger, West Africa | Diversity |
| 2019 | Karmidine Hima | Population Dynamics and Genetics of Gerbillus nigeriae in Central Sahel: Implications for Rodent Pest Control | Ecology and Evolutionary Biology |
| 2019 | Kouame Akpatou | Assessment of Terrestrial Small Mammals in an Agro-industrial Company Concession, Western Liberia | International Journal of Applied Sciences and Biotechnology |
| 2019 | L Karan | Lassa Virus in the Host Rodent Mastomys Natalensis within Urban Areas of Nâ€™zerekore, Guinea | bioRxiv |
| 2019 | Moussa Diagne | Usutu Virus Isolated from Rodents in Senegal | Viruses |
| 2019 | Natalie Weber | New records of bats and terrestrial small mammals from the Seli River in Sierra Leone before the construction of a hydroelectric dam. | Biodiversity Data Journal |
| 2019 | Safianu Rabiu | Demographic response of the Gambian Gerbil to seasonal changes in Savannah fallow fields | Folio Oecologica |
| 2019 | Stefano Catalano | Plagiorchis sp. in small mammals of Senegal and the potential emergence of a zoonotic trematodiasis | IJP: Parasites and Wildlife |
| 2020 | Adama Diarra | Molecular Detection of Microorganisms Associated with Small Mammals and Their Ectoparasites in Mail | American Journal of Tropical Medicine and Hygiene |
| 2020 | Adama Zida | Mastomys natalensis, Cricetomys gambianus and Taterillus sp. were found {PCR} positive for Leishmania major in Burkina Faso, West Africa. | Annals of Parasitology |
| 2020 | Christophe Diagne | Association between temporal patterns in helminth assemblages and successful range expansion of exotic Mus musculus domesticus in Senegal | Biological Invasions |
| 2020 | Claire Stragier | Interplay between historical and current features of the cityscape in shaping the genetic structure of the house mouse ( Mus musculus domesticus) in Dakar (Senegal, West Africa) | Peer Community in Ecology |
| 2020 | Handi Dahmana | Rodents as Hosts of Pathogens and Related Zoonotic Disease Risk | Pathogens |
| 2020 | Henri-Joel Dossou | Invasive rodents and damages to food stocks: a study in the Autonomous Harbor of Cotonou, Benin. | Biotechnologie Agronomie Societe et Environnement |
| 2020 | Joachim Marien | Households as hotspots of Lassa fever? Assessing the spatial distribution of Lassa virus-infected rodents in rural villages of Guinea | Emerging Microbes and Infections |
| 2020 | Laurent Ahissa | Species composition and community structure of terrestrial small mammals in TanoÃ©-Ehy Swamp Forest (South-East Ivory Coast): implication for conservation | Nature Conservation Research |
| 2020 | Leonce Kouadio | Detection of possible spillover of a novel hantavirus in a Natal mastomys from Guinea. | Virus genes |
| 2020 | Stefano Catalano | Multihost Transmission of Schistosoma mansoni in Senegal, 2015-2018 | Emerging Infectious Diseases |
| 2020 | Violaine Nicolas | Small mammal inventory in the Lama forest reserve (south Benin), with new cytogenetical data | Journal of Vertebrate Biology |
| 2021 | Chibuisi Alimba | Wild black rats (Rattus rattus Linnaeus, 1758) as zoomonitor of genotoxicity and systemic toxicity induced by hazardous emissions from Abule Egba unsanitary landfill, Lagos, Nigeria. | Environmental science and pollution research international |
| 2021 | El Hadji Ndiaye | Tick-borne relapsing fever Borreliosis, a major public health problem overlooked in Senegal | PLOS NTD |
| 2021 | Mnqobi Mamba | Small mammals of a West African hotspot, the Ziama-Wonegizi-Wologizi transfrontier forest landscape | Mammalia |
| 2021 | Umaru Bangura | Lassa Virus Circulation in Small Mammal Populations in Bo District, Sierra Leone | Biology |

#### 1.3. Supplementary Table 3

Supplementary Table 3.1: Final GAM model (5). TN_density_ ~ Tweedie(P_density_ + R_area_ + ψ_urban_ + (X * Y)“)

| **Component** | **Term** | **Estimate** | **Std Error** | **t-value** | **p-value** |  |
| --- | --- | --- | --- | --- | --- | --- |
| A. parametric coefficients | (Intercept) | -2.599 | 0.123 | -21.112 | 0.0000 | *** |
| **Component** | **Term** | **edf** | **Ref. df** | **F-value** | **p-value** |  |
| B. smooth terms | s(pop_2005) | 7.131 | 11.000 | 6.398 | 0.0000 | *** |
|  | s(area_km2) | 3.629 | 9.000 | 3.450 | 0.0000 | *** |
|  | s(urban) | 1.921 | 9.000 | 1.235 | 0.0019 | ** |
|  | s(x,y) | 27.252 | 39.000 | 4.563 | 0.0000 | *** |
| Signif. codes: 0 <= '***' < 0.001 < '**' < 0.01 < '*' < 0.05 | | | | | | |
| Adjusted R-squared: 0.301, Deviance explained 0.487 | | | | | | |
| fREML : 3907.563, Scale est: 8.408, N: 1450 | | | | | | |

Supplementary Table 3.2: GAM model 1. TN_density_ ~ Tweedie(X * Y)

| **Component** | **Term** | **Estimate** | **Std Error** | **t-value** | **p-value** |  |
| --- | --- | --- | --- | --- | --- | --- |
| A. parametric coefficients | (Intercept) | -2.062 | 0.126 | -16.403 | 0.0000 | *** |
| **Component** | **Term** | **edf** | **Ref. df** | **F-value** | **p-value** |  |
| B. smooth terms | s(x,y) | 30.431 | 39.000 | 7.290 | 0.0000 | *** |
| Signif. codes: 0 <= '***' < 0.001 < '**' < 0.01 < '*' < 0.05 | | | | | | |
| Adjusted R-squared: 0.0261, Deviance explained 0.349 | | | | | | |
| fREML : 4252.745, Scale est: 11.991, N: 1450 | | | | | | |

Supplementary Table 3.3: GAM model 2. TN_density_ ~ Tweedie(P_density_ + (X * Y))

| **Component** | **Term** | **Estimate** | **Std Error** | **t-value** | **p-value** |  |
| --- | --- | --- | --- | --- | --- | --- |
| A. parametric coefficients | (Intercept) | -2.230 | 0.119 | -18.704 | 0.0000 | *** |
| **Component** | **Term** | **edf** | **Ref. df** | **F-value** | **p-value** |  |
| B. smooth terms | s(pop_2005) | 2.818 | 11.000 | 7.231 | 0.0000 | *** |
|  | s(x,y) | 27.640 | 39.000 | 4.545 | 0.0000 | *** |
| Signif. codes: 0 <= '***' < 0.001 < '**' < 0.01 < '*' < 0.05 | | | | | | |
| Adjusted R-squared: 0.161, Deviance explained 0.422 | | | | | | |
| fREML : 3924.187, Scale est: 9.836, N: 1450 | | | | | | |

Supplementary Table 3.4: GAM model 3. TN_density_ ~ Tweedie(P_density_ + R_area_ + (X * Y))

| **Component** | **Term** | **Estimate** | **Std Error** | **t-value** | **p-value** |  |
| --- | --- | --- | --- | --- | --- | --- |
| A. parametric coefficients | (Intercept) | -2.371 | 0.120 | -19.699 | 0.0000 | *** |
| **Component** | **Term** | **edf** | **Ref. df** | **F-value** | **p-value** |  |
| B. smooth terms | s(pop_2005) | 3.182 | 11.000 | 9.055 | 0.0000 | *** |
|  | s(area_km2) | 3.052 | 9.000 | 2.294 | 0.0000 | *** |
|  | s(x,y) | 26.819 | 39.000 | 4.843 | 0.0000 | *** |
| Signif. codes: 0 <= '***' < 0.001 < '**' < 0.01 < '*' < 0.05 | | | | | | |
| Adjusted R-squared: 0.148, Deviance explained 0.443 | | | | | | |
| fREML : 3941.346, Scale est: 9.432, N: 1450 | | | | | | |

Supplementary Table 3.5: GAM model 4. TN_density_ ~ Tweedie(P_density_ + ψ_tree_ + ψ_urban_ + (X * Y)

| **Component** | **Term** | **Estimate** | **Std Error** | **t-value** | **p-value** |  |
| --- | --- | --- | --- | --- | --- | --- |
| A. parametric coefficients | (Intercept) | -1.775 | 0.123 | -14.379 | 0.0000 | *** |
| **Component** | **Term** | **edf** | **Ref. df** | **F-value** | **p-value** |  |
| B. smooth terms | s(cropland) | 0.000 | 9.000 | 0.000 | 0.7706 |  |
|  | s(shrubland) | 0.000 | 8.000 | 0.000 | 0.4709 |  |
|  | s(tree_cover) | 1.579 | 9.000 | 0.795 | 0.0089 | ** |
|  | s(urban) | 4.772 | 9.000 | 19.703 | 0.0000 | *** |
|  | s(x,y) | 1.837 | 19.000 | 1.499 | 0.0000 | *** |
| Signif. codes: 0 <= '***' < 0.001 < '**' < 0.01 < '*' < 0.05 | | | | | | |
| Adjusted R-squared: 0.0599, Deviance explained 0.306 | | | | | | |
| fREML : 4198.018, Scale est: 12.895, N: 1450 | | | | | | |

#### 1.4. Supplementary Fig 1

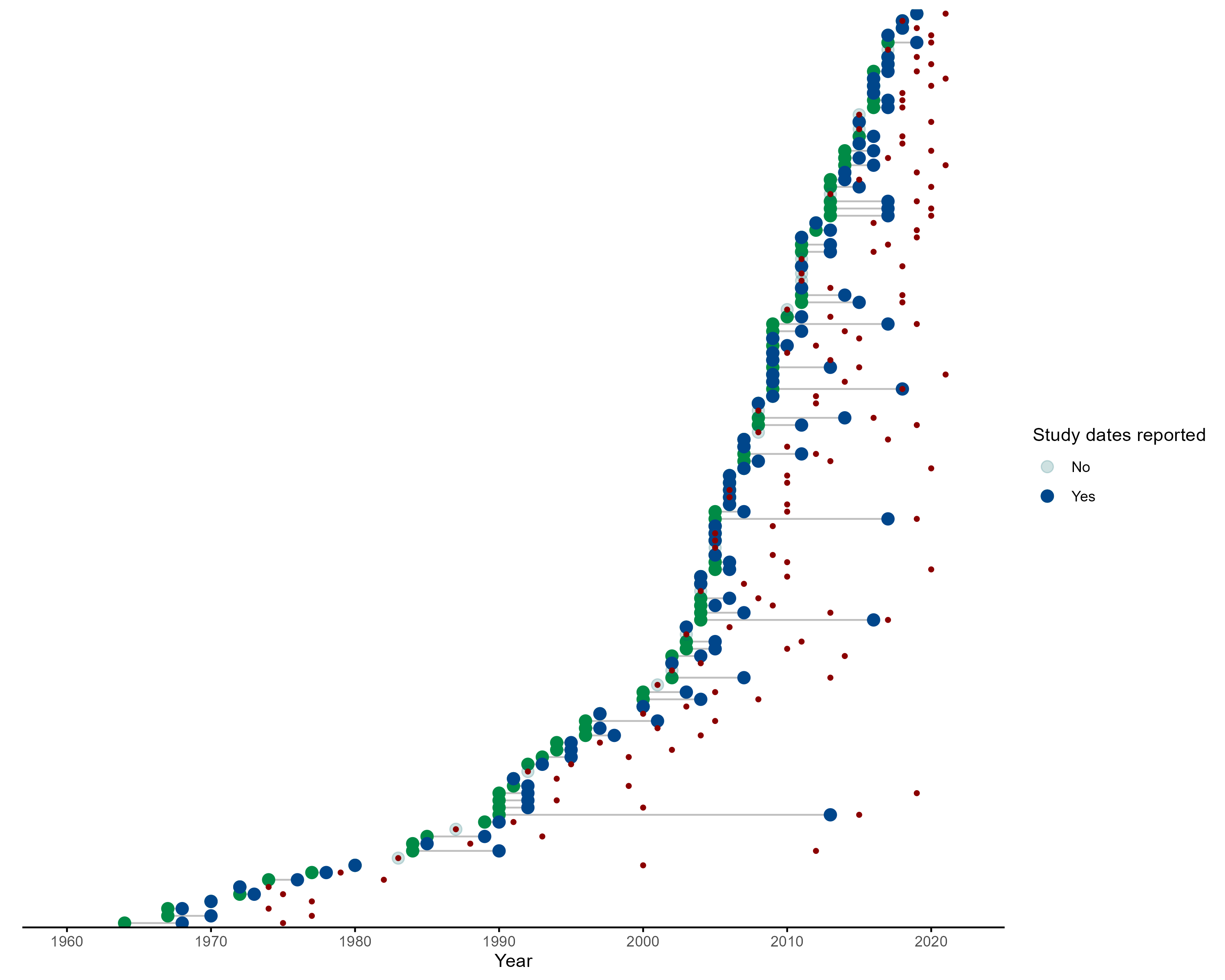

Supplementary Fig 1. Green points represent the start date of rodent trapping studies, blue points representing the final trapping activity. Red points indicate the publication of studies. Increasing numbers of studies have been published since 2000 with more studies being conducted over repeated visits.

#### 1.5. Supplementary Fig 2

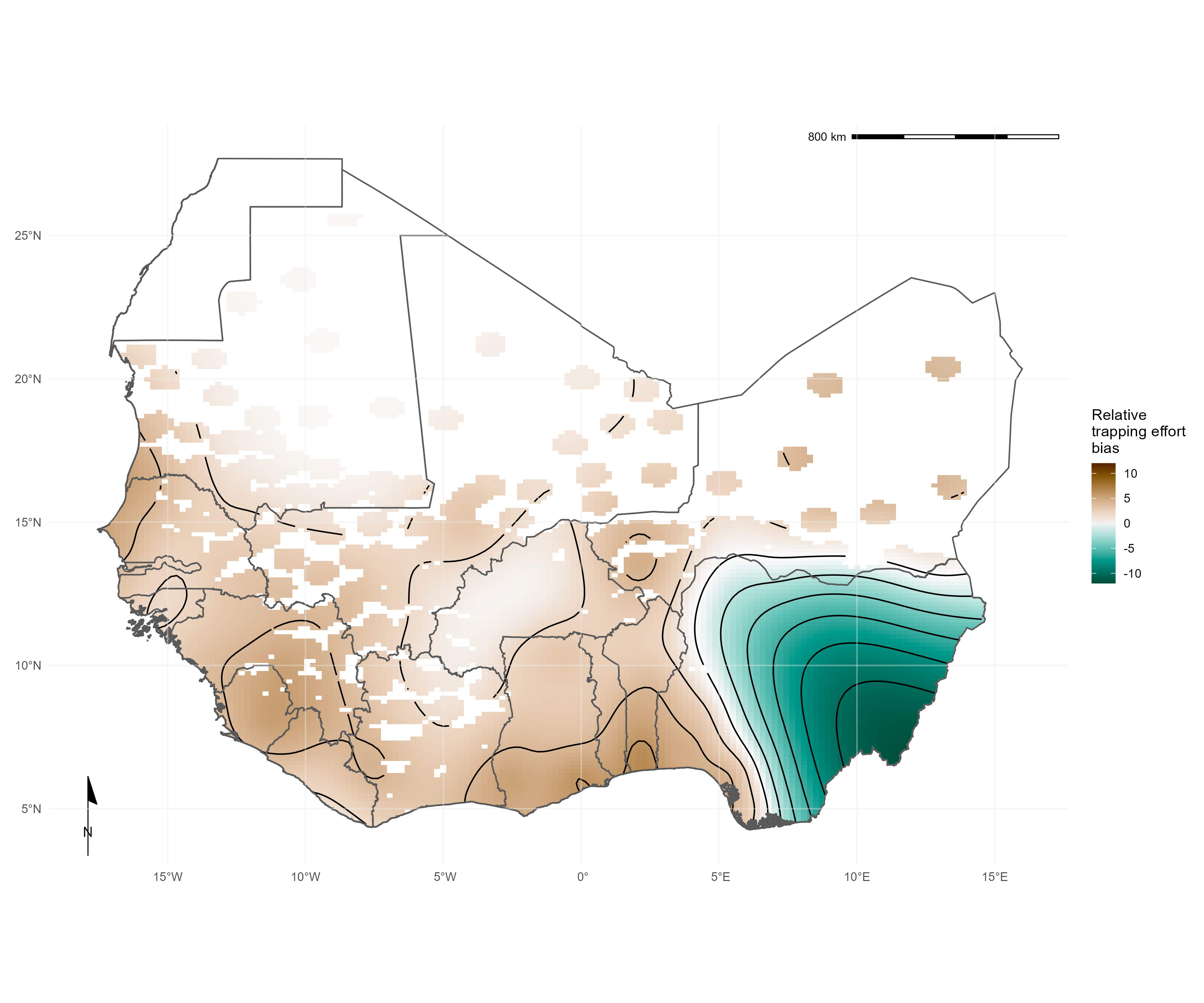

Supplementary Fig 2. Relative trapping effort bias across West Africa from the subset of included studies reporting trapping effort, adjusted for proportion urban land cover and proportion tree cover. Brown regions represent areas with higher than expected trapping effort, green regions represent areas lower than expected trapping effort

#### 1.6. Supplementary Fig 3

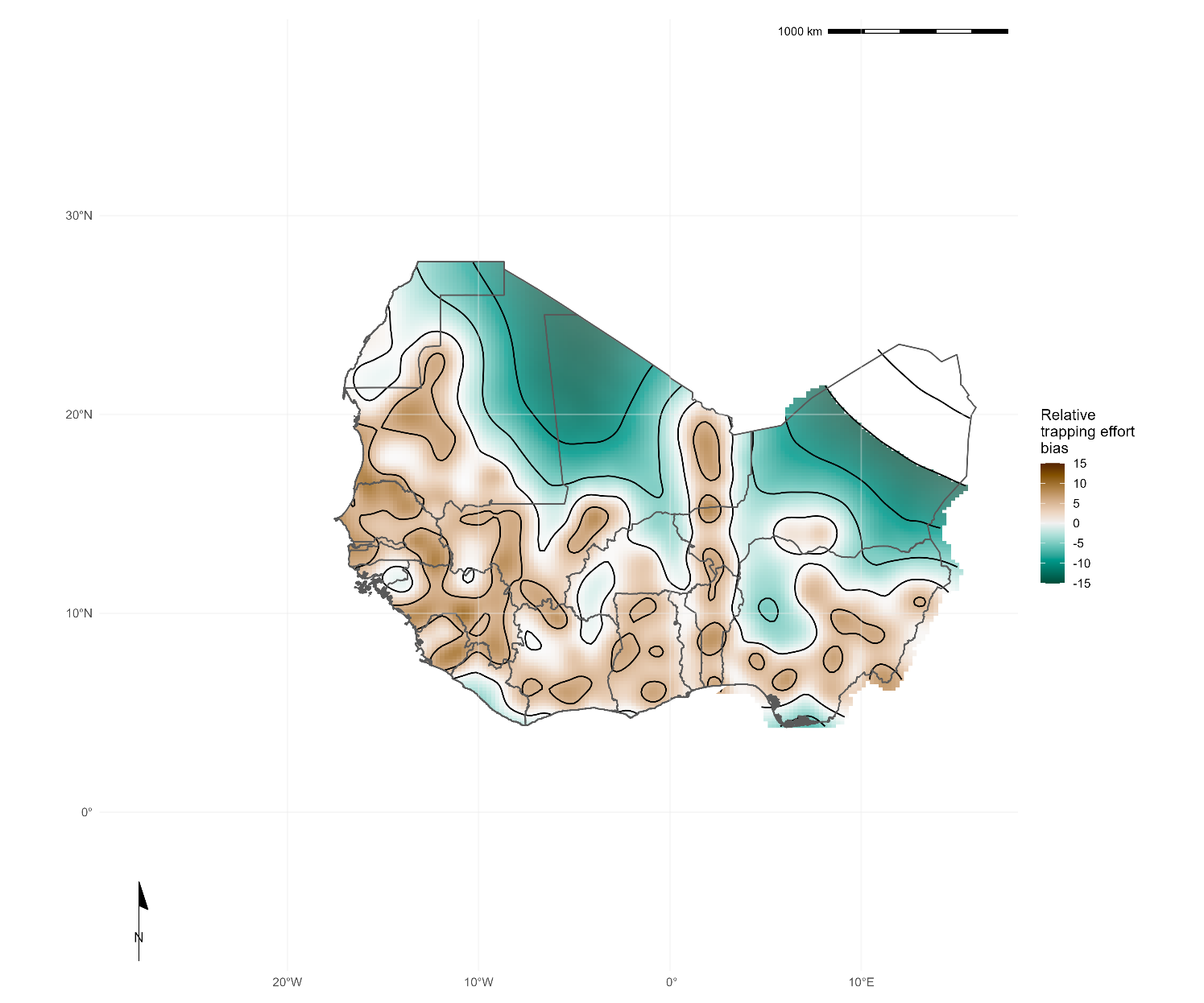

Supplementary Fig 3. Pixel based analysis of relative trapping effort bias across West Africa adjusted for habitat type and human population density. Brown regions represent areas with higher than expected trapping effort, green regions represent areas lower than expected trapping effort

#### 1.7. Supplementary Fig 4

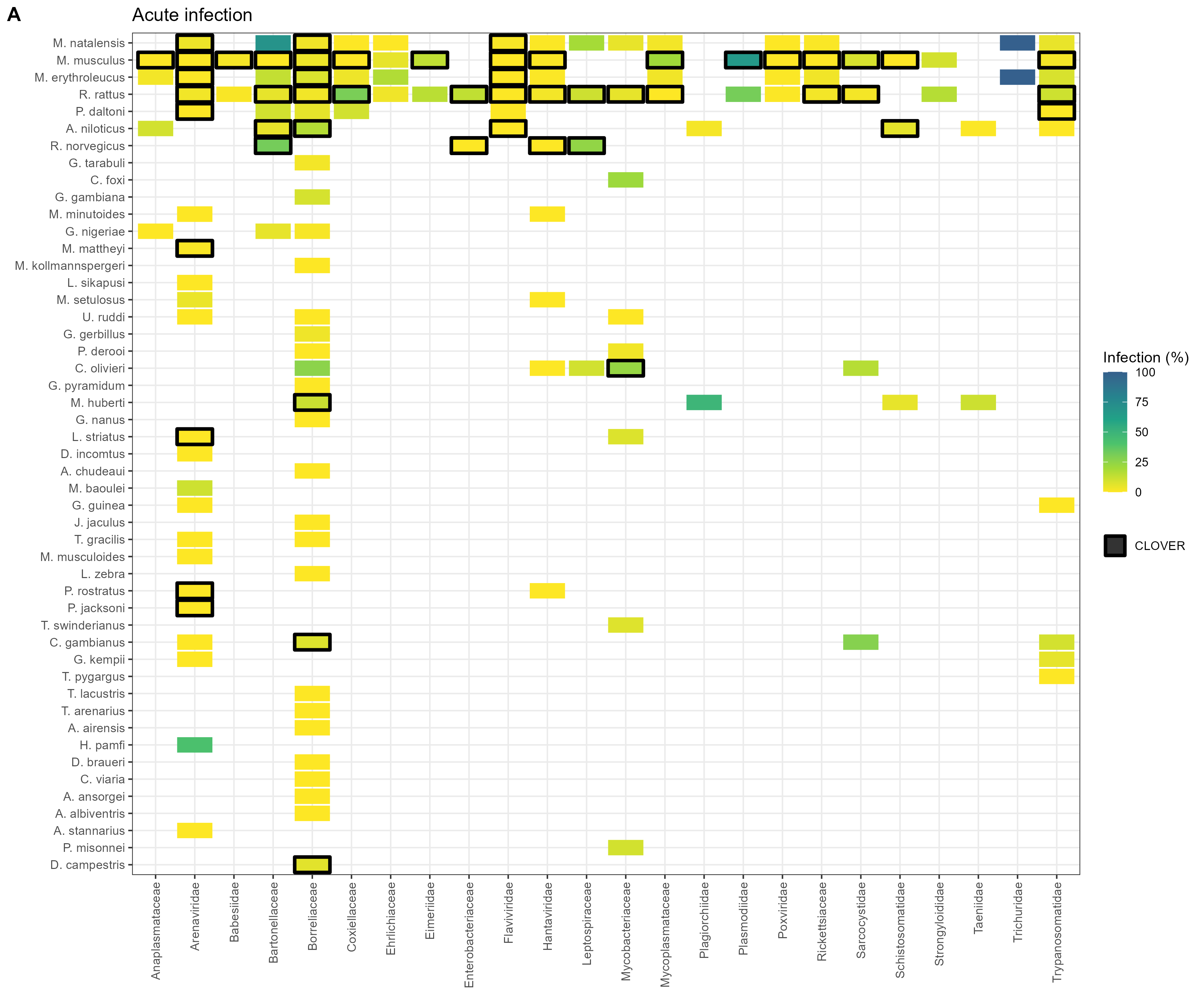

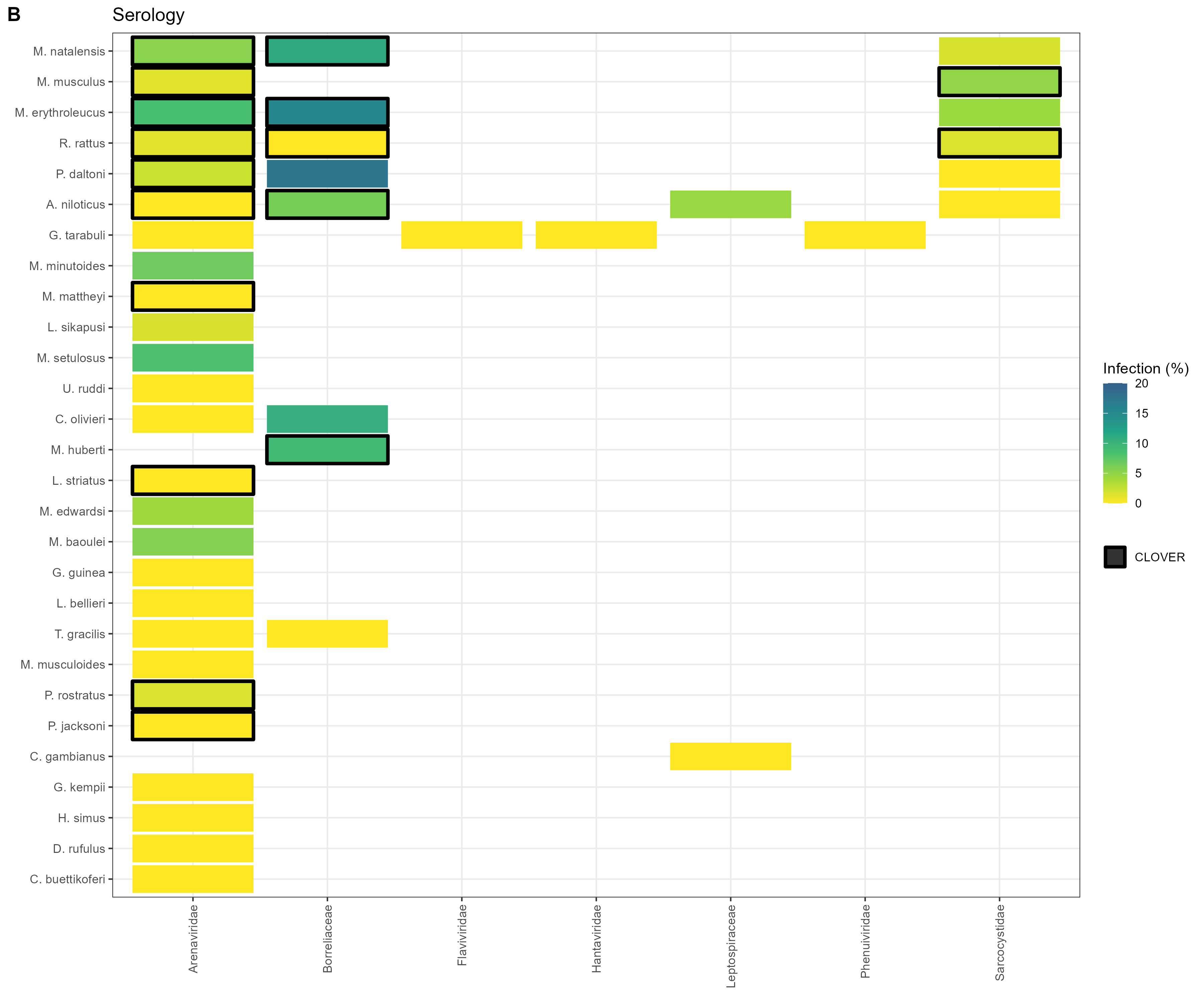

Supplementary Fig 4. A) Identified host-pathogen associations at pathogen family level through detection of acute infection (i.e. PCR, culture). Percentages and colour relate to the proportion of all assays that were positive. Associations with a black border are present in the CLOVER dataset. B) Identified host-pathogen associations at pathogen family level through serological assays (i.e. ELISA). Percentages and colour relate to the proportion of all assays that were positive. Associations with a black border are present in the CLOVER dataset.

#### 1.8. Supplementary Fig 5

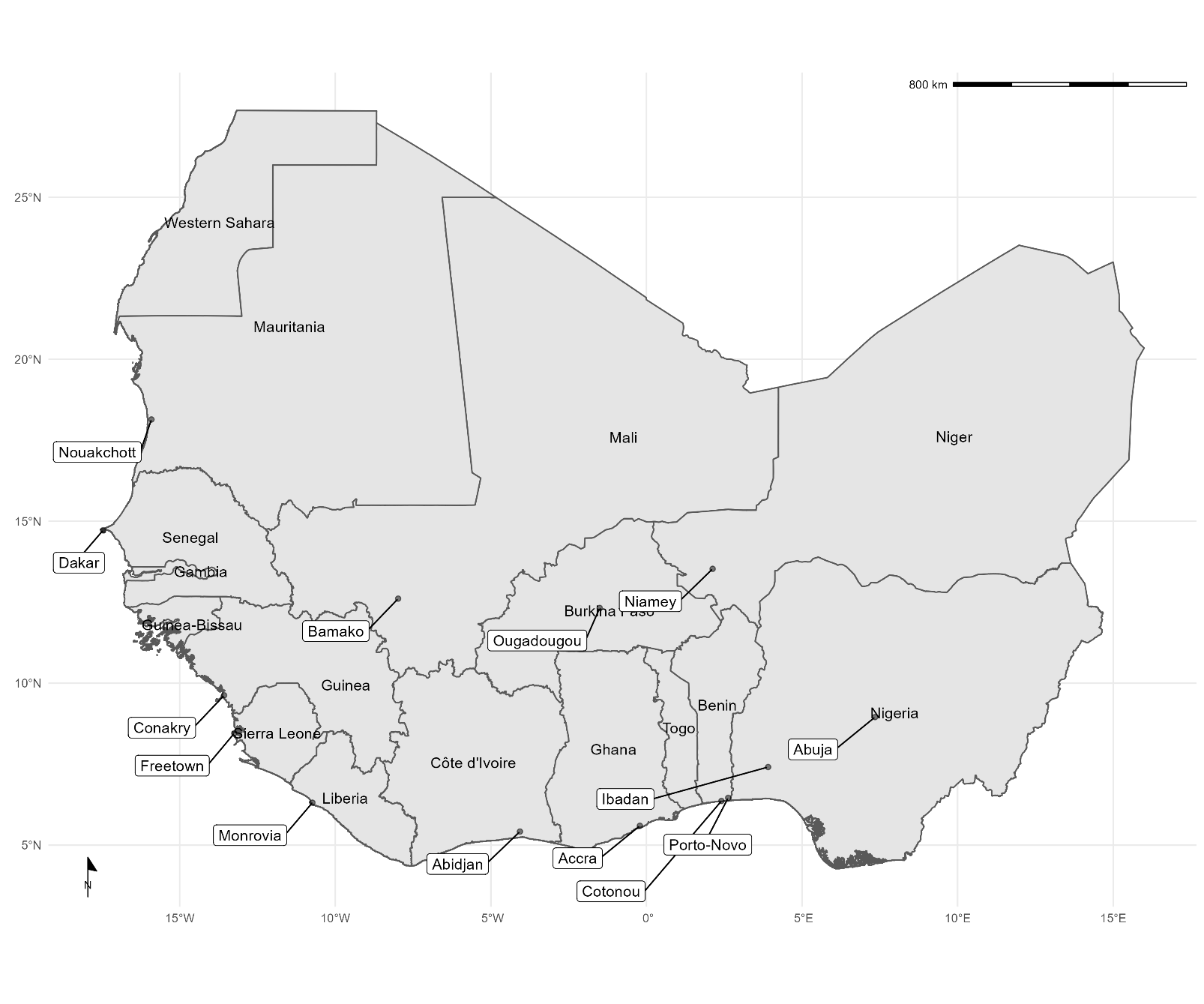

Supplementary Fig 5. A map of the study region with capital cities and areas discussed in the manuscript highlighted.

#### 1.9. Supplementary Fig 6

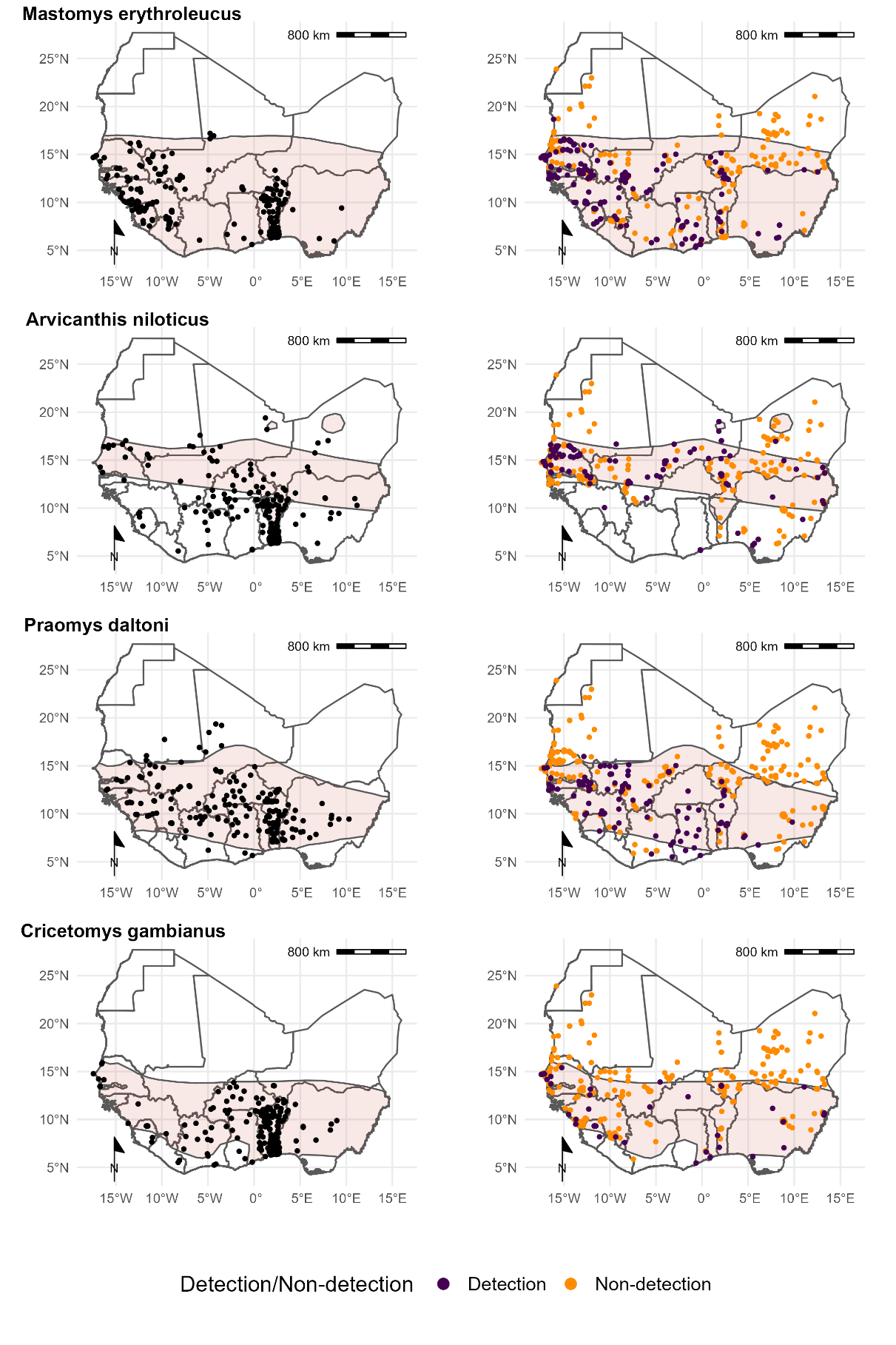

Supplementary Fig 6. Locations of detection and non-detection sites for rodent species in West Africa. Each row corresponds to a single rodent species. L) Presence recorded in GBIF (black points) overlaid on IUCN species range (red-shaded area). R) Detection (purple) and non-detection (orange) from rodent trapping studies overlaid on IUCN species ranges. M. musculus has no IUCN West African range.
